# E3 Ligase Ligandalyzer: A Structure-First, Ligand-Centric Platform for E3 Ligase Recruiter Selection

**DOI:** 10.64898/2026.10.07.757351

**Authors:** Joseph M. Schulz, Mainak Bardhan, Robert C. Reynolds, Gary LeRoy, Danny Reinberg, Stephan C. Schürer

**Affiliations:** Department of Molecular and Cellular Pharmacology, University of Miami, Miami, FL, USA; Howard Hughes Medical Institute, University of Miami, Miami, FL 33136, USA; The Dr. John T. Macdonald Foundation Department of Human Genetics, University of Miami Miller School of Medicine, Miami, FL, U.S.A.; The John P. Hussman Institute for Human Genomics, University of Miami Miller School of Medicine, Miami, FL, U.S.A; Sylvester Comprehensive Cancer Center, University of Miami Miller School of Medicine, Miami, FL, USA; O’Neal Comprehensive Cancer Center, University of Alabama at Birmingham, Birmingham, AL 35205, USA; Frost Institute for Data Science & Computing, University of Miami, Miami, FL, USA

**Keywords:** E3 ubiquitin ligase, targeted protein degradation, PROTAC, recruiter ligand, structural database

## Abstract

Here, we present E3 Ligase Ligandalyzer Version 1.0, a freely accessible, recruiter-centric web platform for comparative exploration of experimentally resolved E3 ligase-recruiter complexes. The locked Version 1.0 dataset spans 35 E3 ligases, 604 reconciled recruiter entities, and 1,372 active structure-specific recruiter instances across 685 experimentally resolved PDB structures. Recruiters are linked to normalized chemical representations and analyzed using cheminformatics and structure-based workflows to compute physicochemical descriptors, scaffold relationships, aligned binding-pocket views, and solvent-accessible surface area patterns relevant to linker attachment. The platform integrates recruiter-entity, recruiter-instance, scaffold, and ligase views, allowing users to compare experimentally grounded chemical and structural observations without imposing predictive rankings. The platform supports direct transition into downstream degrader-design workflows, including integration with PROTAC Builder (protacbuilder.com). E3 Ligase Ligandalyzer is freely available at https://e3ligandalyzer.com, and the associated code and dataset-generation scripts are available at https://github.com/schurerlab/E3-Ligandalyzer-Scripts.

Database URL: https://e3ligandalyzer.com

## Introduction

Targeted protein degradation (TPD) enables the elimination of disease-relevant proteins rather than requiring inhibition of a single functional site (1). Technologies such as PROTACs, molecular glues, and related degrader modalities exploit the ubiquitin-proteasome system by redirecting E3 ubiquitin ligases toward selected protein targets (2). Despite the rapid growth of the field, the rational design of degrader molecules remains challenging, particularly at early discovery stages (3,4).

A central early decision in TPD development is the selection of an E3 ligase recruiter. While hundreds of E3 ligases exist in the human genome, only a limited subset has been routinely exploited for drug discovery (5,6). Even among commonly used ligases, such as CRBN, VHL, IAPs, and MDM2, recruiter ligands vary in chemical tractability, physicochemical properties, binding modes, affinity, and available positions for linker attachment (7). The early development of tractable recruiter systems, including VHL-binding ligands, helped concentrate design around a small number of established chemotypes (8). Recruiter selection is therefore often guided by historical precedent rather than direct comparison, even though the choice constrains the chemical properties and geometries that can be explored in a degrader series (9,10).

Although structural biology, cheminformatics, and docking approaches have been widely applied to degrader design, information needed to compare E3 recruiters remains distributed across PDB entries, publications, and specialized databases. Resources centered on complete degrader molecules and associated activities include PROTAC-DB and its updated releases, TPDdb, and PROTAC-Databank (11–15). Other resources organize E3 biology and substrate relationships (5,16–19). These resources are organized around those respective entities rather than a common representation for direct comparison of experimentally resolved recruiter-ligase complexes. Researchers must therefore reconcile ligand identifiers, chemical structures, bound conformations, and biological annotations before comparing recruiters across ligases. This fragmentation is particularly important at early design stages, where molecular size, polarity, ligand efficiency, binding geometry, and linker compatibility must be considered together. These considerations reflect a broader principle of targeted conjugate design: molecular scale, linker architecture, cellular access, and whole-molecule physicochemical behavior must be optimized together because gains in selective recognition can be offset by limitations in exposure, trafficking, or productive pharmacology (20).

Several peer-reviewed resources address complementary aspects of TPD. PROTAC-DB aggregates structures and activity data for complete PROTACs, including recruiter and linker components (11–13). TPDdb consolidates degraders across modalities with chemical and biological annotations (14). ELiAH and the E3 Ligase Atlas provide tissue- and cell-type-specific expression context for human E3 ligases (6,21). ELIOT maps ligandable pockets across E3 ligases (22). These resources support degrader cataloging, ligase selection, or recruiter discovery, but their primary records are not organized for direct cross-ligase comparison of experimentally resolved recruiter chemotypes.

Complementary computational tools focus on assembled degraders or ternary-complex modeling. PROTACable links recruiter and warhead fragments to generate candidate degraders (23). PRosettaC, DegraderTCM, and MEGA PROTAC sample or rank ternary complexes (24–26). PRosettaC has also been benchmarked against AlphaFold 3 for this task (27). DeepPROTACs, DiffPROTACs, DeepPSA, and AI-DPAPT predict degradation-related outcomes, generate candidate molecules, or estimate synthetic accessibility (28–31). Physics-based and integrative approaches, including SILCS-xTAC, induced-fit docking with molecular dynamics, and IMP-based modeling, examine predefined degrader components or assembled systems (32–34). A recent computational Perspective organizes these approaches into complementary constraint-driven, physics-based, and data-driven workflows while highlighting persistent limitations in benchmarking, scoring, molecular representation, and prospective deployment (35). PROTAC-PatentDB provides a complementary patent-derived collection of complete degraders (36). P4ward provides a complementary, fully automated and open-source workflow for PROTAC ternary-complex modeling from defined binary complex structures and PROTAC inputs, combining protein-protein docking, linker sampling, filtering, and scoring to support the screening of multiple candidate degraders (37). These methods address downstream design questions but generally require recruiter identities, attachment sites, or structural inputs to be supplied.

Recent methods expand this downstream landscape while retaining a dependence on well-defined structural or chemical inputs. The PROTAC Conformer Generator (PCG) preserves protein-bound E3-recruiting-group and warhead geometries during linker sampling and rigid-body ternary-complex assembly (38). DeepTernary uses an SE(3)-equivariant deep-learning framework for ternary-complex prediction (39), whereas TERNIFY restricts sampling to linker conformational degrees of freedom (40) and COMPASS screens linker libraries for ternary-complex feasibility and ubiquitination potential (41).

The absence of a unified, recruiter-centric resource is particularly limiting for constraint-driven, geometry-based degrader design strategies (9). Such methods rely on accurate representations of recruiter binding modes, attachment directions, and linker constraints to generate physically plausible ternary-complex models (10,42–44). Experimentally determined recruiter–ligase complexes provide these observed binding contexts. Solvent exposure and bound geometry can help identify positions for further investigation, but they do not establish linker tolerance or productive degradation.

**Table 1** summarizes representative degrader-centric databases and computational tools. These resources span curated degrader datasets, data-driven predictors, and structure-based modeling methods, but they address a different unit or stage of analysis from recruiter-level structural curation.

**Table 1.** Representative degrader-centric databases and computational tools for PROTAC design and modeling.

| Resource/tool | Focus/description | DOI |
| --- | --- | --- |
| **PROTAC-DB** | A curated database of PROTAC structures, warheads, recruiters and linkers with activity data. Provides search and download functions for degrader molecules and their components. | 10.1093/nar/gkaa807 |
| **PROTACable** | Integrative pipeline combining 3-D modeling and deep learning to automate de novo PROTAC design and scoring. Generates candidate degraders by linking recruiter and warhead fragments and ranks them using predicted ternary complex stability. | 10.1021/acs.jcim.3c01878 |
| **PRosettaC** | Rosetta-based protocol for modeling PROTAC-mediated ternary complexes. Uses docking and scoring routines to sample recruiter–target–ligase configurations and predict favourable ternary geometries. | 10.1021/acs.jcim.0c00589 |
| **DegraderTCM** | Computationally efficient method for predicting PROTAC ternary complexes using a combination of docking, resampling and scoring. Designed to work even when structural data are sparse. | 10.1021/acsmedchemlett.3c00362 |
| **MEGA PROTAC (MEGADOCK)** | Pipeline that uses high-throughput protein–protein docking (MEGADOCK) followed by sequential filtering and rank aggregation to model ternary complexes and prioritise PROTAC designs. | 10.1038/s41598-024-83558-2 |
| **P4ward** | Fully automated, open-source pipeline for PROTAC ternary-complex modeling from two binary protein–ligand complexes and one or more PROTAC structures. Integrates protein–protein docking, linker sampling, filtering, and scoring to generate ranked ternary-complex models and support screening across multiple degraders. | 10.1021/acs.jcim.5c00614 |
| **PROTAC Conformer Generator (PCG)** | Knowledge-based workflow for PROTAC conformer generation and rigid-body ternary-complex assembly that preserves protein-bound E3-recruiter and warhead geometries during linker sampling. | 10.1021/acs.jcim.5c00880 |
| **COMPASS** | Structure-based pipeline that screens PROTAC linker libraries for ternary-complex feasibility and ubiquitination potential, supporting elimination of nonproductive candidates before synthesis. | 10.1002/cmdc.70385 |
| **Induced-fit docking & MD simulation of VHL-mediated PROTACs** | Case study modelling ternary complexes of VHL recruiter and FLT3 warhead using induced-fit docking and molecular dynamics simulations; illustrates the importance of geometry and dynamics in degrader design. | 10.1002/ardp.202500102 |
| **DeepPROTACs** | Deep learning model that predicts degradation outcomes (e.g., ternary complex formation and degradation potency) from PROTAC structures. Incorporates graph neural networks and attention mechanisms. | 10.1038/s41467-022-34807-3 |
| **DiffPROTACs** | Diffusion-based generative model that designs new PROTACs from scratch. Trains on known degraders and uses a diffusion process to propose candidate molecules. | 10.1093/bib/bbae358 |
| **DeepPSA** | Geometric deep-learning approach for predicting synthetic accessibility (how easily a PROTAC can be synthesised) using protein and ligand structures. | 10.1021/acs.jcim.5c00366 |
| **AI-DPAPT** | Machine-learning framework for predicting PROTAC activity by combining molecular descriptors of warheads, linkers and recruiters. Provides predictions of degradation efficacy for new degrader designs. | 10.1007/s11030-024-11011-7 |
| **SILCS-xTAC** | An extension of the SILCS (Site Identification by Ligand Competitive Saturation) method for modeling PROTAC ternary complexes. Generates ensembles of ternary complexes and calculates binding free energies using fragment affinity maps. | 10.1021/acs.jcim.5c02045 |
| **PRosettaC vs AlphaFold-3 benchmarking** | Scientific Reports study comparing PRosettaC with AlphaFold-3 for modeling PROTAC-mediated ternary complexes; shows that constraint-driven approaches still outperform end-to-end predictions in most cases. | 10.1038/s41598-025-21502-8 |
| **Integrative modeling of PROTAC ternary complexes** | Integrative Modeling Platform (IMP) workflow for assembling ternary complexes using experimental restraints (e.g., crosslinking). Provides scoring functions and sampling algorithms for PROTAC design. | 10.1021/acs.jmedchem.1c01576 |
| **PROTAC-PatentDB** | Dataset of PROTAC compounds extracted from patent literature (2013–2023) with associated chemical structures and metadata. Useful for screening novel recruiter–warhead combinations. | 10.1038/s41597-025-06136-9 |

**Table 2** summarizes representative resources addressing the biological and structural properties of E3 ligases. These atlases, interaction networks, ubiquitination databases, and pocket-mapping tools provide expression profiles, substrate interactions, evolutionary relationships, or ligandability information, rather than a common set of recruiter-complex records.

**Table 2.**
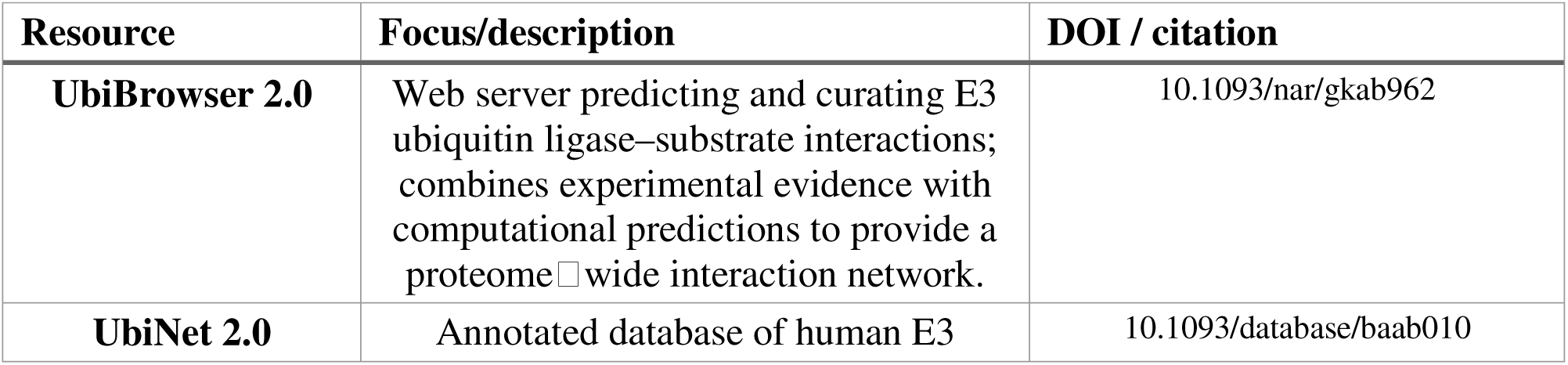
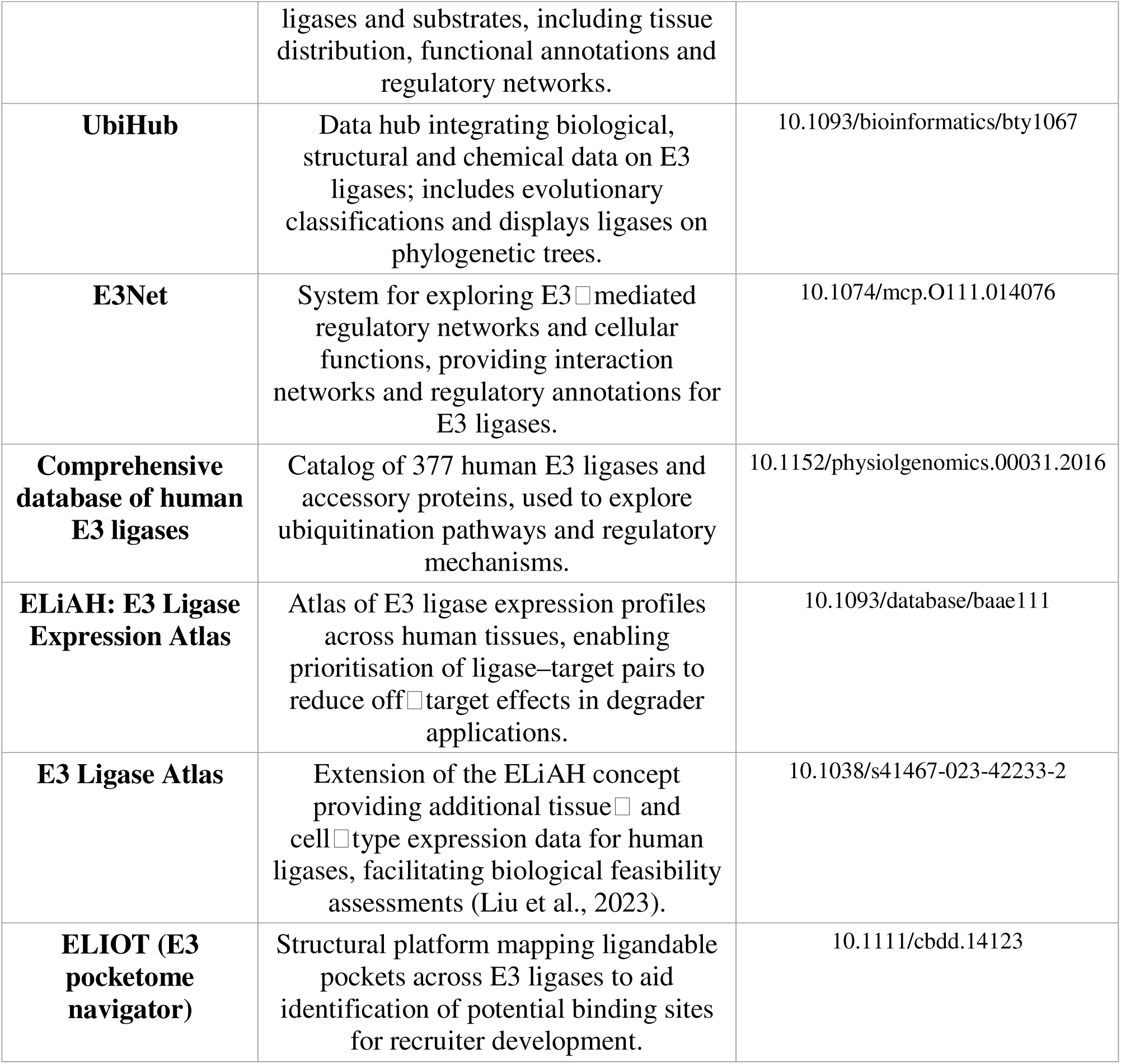
Representative E3 ligase-centered databases and resources.

Together, the resources in **Tables 1** and **2** cover complete degrader chemistry, E3 ligase biology, expression, and pocket identification (6,13,21,22). What remains difficult is direct comparison of experimentally resolved recruiter chemotypes across ligases in a consistent chemical and structural representation. Such comparison requires linking ligand/component identifiers to specific structures, retaining repeated binding contexts, standardizing chemical records, grouping related scaffolds, and calculating comparable structural annotations.

To address this need, we developed E3 Ligase Ligandalyzer, an interactive, web-based database that catalogs and compares experimentally resolved E3 ligase-recruiter complexes. The locked Version 1.0 dataset, with a database cutoff of 8 September 2026, contains 604 reconciled recruiter entities represented by 1,372 active recruiter instances across 685 PDB structures and 35 curated E3 ligases. A Recruiter_ID denotes the reconciled chemical recruiter entity, whereas a Recruiter_Instance_ID denotes an exact, structure-specific observation. Thus, one recruiter entity may be retained as multiple instances when it occurs in distinct experimentally resolved structures or binding contexts.

Recruiter entities are linked to calculated physicochemical descriptors, Bemis-Murcko scaffold assignments, and aligned binding-site views. Recruiter instances retain atom-level SASA annotations and experimentally observed structural coordinates for each bound complex. The active Version 1.0 scientific population contains 428 scaffolds and supports recruiter-entity, recruiter-instance, scaffold, and ligase-centered comparison without treating structural observations as predictions of degrader activity.

Selected biological annotations derived from ELiAH are incorporated to provide expression context alongside the structural and chemical records (21). Expression data are not used to rank ligases or recruiter compounds. Atom-level SASA can highlight positions that remain exposed in a bound complex, but it is presented as a descriptive annotation rather than a validated predictor of linker attachment.

Users can explore recruiter ligands across multiple E3 ligases, compare design-relevant properties, and visually inspect experimentally determined binding modes. Selected recruiter records can also be exported to PROTAC Builder for optional downstream molecule construction, while E3 Ligase Ligandalyzer remains usable as a standalone database.

This manuscript describes the curation strategy, database organization, and representative analyses of E3 Ligase Ligandalyzer. The resource is intended to support comparison and hypothesis generation at the recruiter-selection stage, not to rank E3 ligases, predict degradation, or validate a linker attachment site. The public interface is available at https://e3ligandalyzer.com, and the dataset-generation scripts are available at https://github.com/schurerlab/E3-Ligandalyzer-Scripts.

## Materials and Methods

### Structural data collection and curation of E3 recruiter complexes

We surveyed RCSB Protein Data Bank (PDB) entries associated with curated human E3 ligases to identify experimentally resolved recruiter-bound structures. Candidate entries were assessed with ligase-specific structural collection and curation workflows, and recruiter observations were retained with their PDB, model, asymmetry, residue, and source-instance provenance. The resulting collection includes conventional chemical-component observations and deposited BIRD/PRD recruiter entities; it is not restricted to conventional small-molecule HETATM records.

For reproducibility, all quantitative analyses in this manuscript use the locked Version 1.0 dataset, with a database cutoff of 8 September 2026. PDB structures released after this cutoff are not included in the manuscript population.

Each accepted observation was assigned a permanent Recruiter_Instance_ID and retained as a structure-specific record. Reconciliation groups chemically equivalent observations under a Recruiter_ID while preserving the associated PDB entry, model, asymmetry identifier, residue identifier, ligase, and observed coordinate provenance. Active records comprise the Version 1.0 scientific population; historical inactive registry rows are preserved for provenance but are excluded from manuscript statistics. This process yielded 1,372 active recruiter instances across 35 E3 ligases.

### Ligand extraction, standardization, and scaffold analysis

For each active recruiter instance, the pipeline generated or selected a normalized 2D chemistry asset while preserving the experimentally observed PDB coordinates as the authority for the 3D structural view. Chemistry provenance was recorded as EXACT_INSTANCE_SDF when a validated instance-specific SDF was available, CANONICAL_SDF when reconciled canonical chemistry supported the observation, or DEPOSITED_CIF_DERIVED when deposited chemical-component definitions were used. BIRD/PRD observations were identified before inappropriate small-molecule filtering and preserved as whole recruiter graphs rather than fragmented residue-level records. These 2D chemistry assets support descriptors and atom correspondence; they do not replace the observed three-dimensional coordinates.

Core scaffolds were derived from normalized recruiter-entity chemistry using Bemis-Murcko decomposition. Entity-level scaffold assignments support grouping of chemical identities across ligases, whereas structure-specific instances remain available for observed-pose and SASA analyses. For each ligase, scaffold Shannon diversity was calculated from recruiter-entity counts distributed across its scaffold assignments as H = -sum(p_i ln p_i); normalized diversity was H divided by ln(S), where S is the number of distinct scaffolds for that ligase. Recruiter-entity density is the recruiter-entity count assigned to a scaffold divided by the total recruiter-entity count for the ligase. The active Version 1.0 scientific population contains 428 scaffold rows; a 429th registry row is historical inactive and is excluded from manuscript-facing statistics.

### Physicochemical and topological descriptor calculation

A panel of physicochemical and topological descriptors relevant to degrader design was computed from reconciled recruiter-entity chemistry using RDKit. These include molecular weight, calculated lipophilicity, topological polar surface area, hydrogen-bond donor and acceptor counts, ring counts, fraction Csp3, molecular refractivity, quantitative estimate of drug-likeness, and topological complexity indices. Entity-level calculation prevents repeated structural observations of the same chemistry from disproportionately influencing intrinsic chemical-property summaries.

All descriptor calculations were performed using Python scripts to ensure consistency across the dataset and facilitate reproducibility. Descriptor values are stored alongside structural metadata and scaffold annotations within the database.

### Structure-based binding pocket analysis and solvent accessibility

Recruiter ligands were analyzed within the context of their experimentally resolved binding sites to enable structure-informed comparison across E3 ligases. Protein–ligand complexes were aligned at the binding pocket level to facilitate qualitative inspection of ligand orientation, exit vector directionality, and pocket geometry across ligases.

SASA was calculated with the Shrake-Rupley algorithm using a 1.4 Å rolling probe. The analytical SASA and 3D-to-2D mapping domains include heavy atoms only: explicit H, D, and T records are excluded from these derived analytical views without mutating the deposited source coordinates. The Version 1.0 validation set contains 41,618 analytical heavy atoms, all mapped to the normalized chemistry representation, with no unresolved heavy atoms and no analytical H/D/T records. SASA is presented as a descriptive structural annotation, not as a validated predictor of linker tolerance or productive degradation.

### Integration of expression-based annotations

To provide biological context without duplicating existing expression resources, selected tissue-level expression annotations were incorporated from published E3 ligase expression atlases (21). These data were integrated without modification and are used solely for visualization and contextual guidance. Expression data are not used to rank, filter, or score recruiter ligands, and the platform’s primary analytical emphasis remains on structure- and ligand-level properties relevant to degrader design.

### Web platform architecture and implementation

E3 Ligase Ligandalyzer is implemented as a web-based application built using the Flask framework and deployed on a Linux server environment (e3ligandalyzer.com). The relational data model separates reconciled recruiter entities from structure-specific recruiter instances. Normalized 2D chemical assets, descriptor data, scaffold assignments, and observed PDB-coordinate assets are linked through recorded provenance, enabling cross-reference between E3 ligases, recruiter chemistry, scaffolds, and structural observations.

The front end supports interactive exploration through dynamic visualizations and embedded molecular viewers. The two-dimensional view displays normalized recruiter chemistry and atom-level SASA correspondence; the three-dimensional viewer displays experimentally observed instance coordinates in their structural context. Interactive plots and network representations support exploration of scaffold relationships, recruiter-entity associations across ligases, and structure-informed design features.

### Interactive exploration and decision-support functionality

The platform is designed as a decision-support tool rather than a static repository. Users can explore recruiter ligands across E3 ligases, compare scaffold usage and physicochemical properties, visualize aligned binding pockets, and examine solvent exposure patterns relevant to linker attachment and optimization. Network-based visualizations enable identification of shared scaffolds and chemotypes across ligases, supporting hypothesis generation for recruiter repurposing and cross-ligase design strategies.

E3 Ligase Ligandalyzer is implemented as a standalone Flask-based web application and integrates directly with PROTAC Builder for downstream degrader development. Recruiter ligands selected within E3 Ligase Ligandalyzer can be transferred directly into the PROTAC Builder environment, where they are loaded as editable two-dimensional chemical structures within an embedded sketch editor. This enables users to modify recruiter chemistry, define attachment points, and construct linker connections interactively, without requiring external cheminformatics software or manual file conversion.

PROTAC Builder further integrates recruiter inputs from E3 Ligase Ligandalyzer with warhead selection tools developed in parallel, including VLiSEMOD (vlisemod.com) for viral target and ligand development. Together, these standalone yet interoperable tools support a modular, constraint-driven PROTAC design workflow in which warhead, recruiter, linker, and ternary complex selection are evaluated within a unified, structure-informed framework.

### Availability

E3 Ligase Ligandalyzer is freely accessible at https://e3ligandalyzer.com. The platform enables interactive exploration of curated recruiter ligands, structures, and scaffold relationships, and this manuscript presents representative analyses to illustrate its scope and utility.

## Results

### Platform content and dataset overview

E3 Ligase Ligandalyzer is a structure-centric web platform for comparative analysis of experimentally resolved E3 ligase-recruiter complexes. The locked Version 1.0 dataset contains 604 reconciled recruiter entities represented by 1,372 active structure-specific recruiter instances across 685 unique PDB structures and 35 E3 ligases. A recruiter entity is a reconciled chemical identity, whereas a recruiter instance is an exact observed occurrence in a specific structure and binding context. Retaining instances separately preserves structural provenance, observed binding modes, pocket context, and linker-relevant geometry without describing all observations as unique ligands.

Recruiter entities were grouped into scaffold classes based on shared chemical backbones, enabling recruiter-entity, recruiter-instance, ligase, and scaffold-centered analysis. The active scientific population contains 428 scaffolds. There are 452 ligase-scaffold associations and 617 ligase-recruiter associations, providing a basis for evaluating scaffold reuse and chemical convergence across E3 ligases.

Summary statistics on the platform describe entity-level recruiter chemistry and association-level scaffold coverage (Figure 1). Across the 604 reconciled recruiter entities, the median Bertz complexity is 1036.1, mean QED is 0.55, and 84.9% satisfy the recorded Lipinski rule-of-five criterion. Across ligases, the mean scaffold Shannon diversity is 1.505 and mean normalized scaffold diversity is 0.653, calculated from recruiter-entity counts per scaffold within each ligase. Structure-specific mapping validation comprises 41,618 analytical heavy atoms and 41,618 mapped heavy atoms, with no unresolved atoms and no analytical H/D/T records.

**Figure 1.**
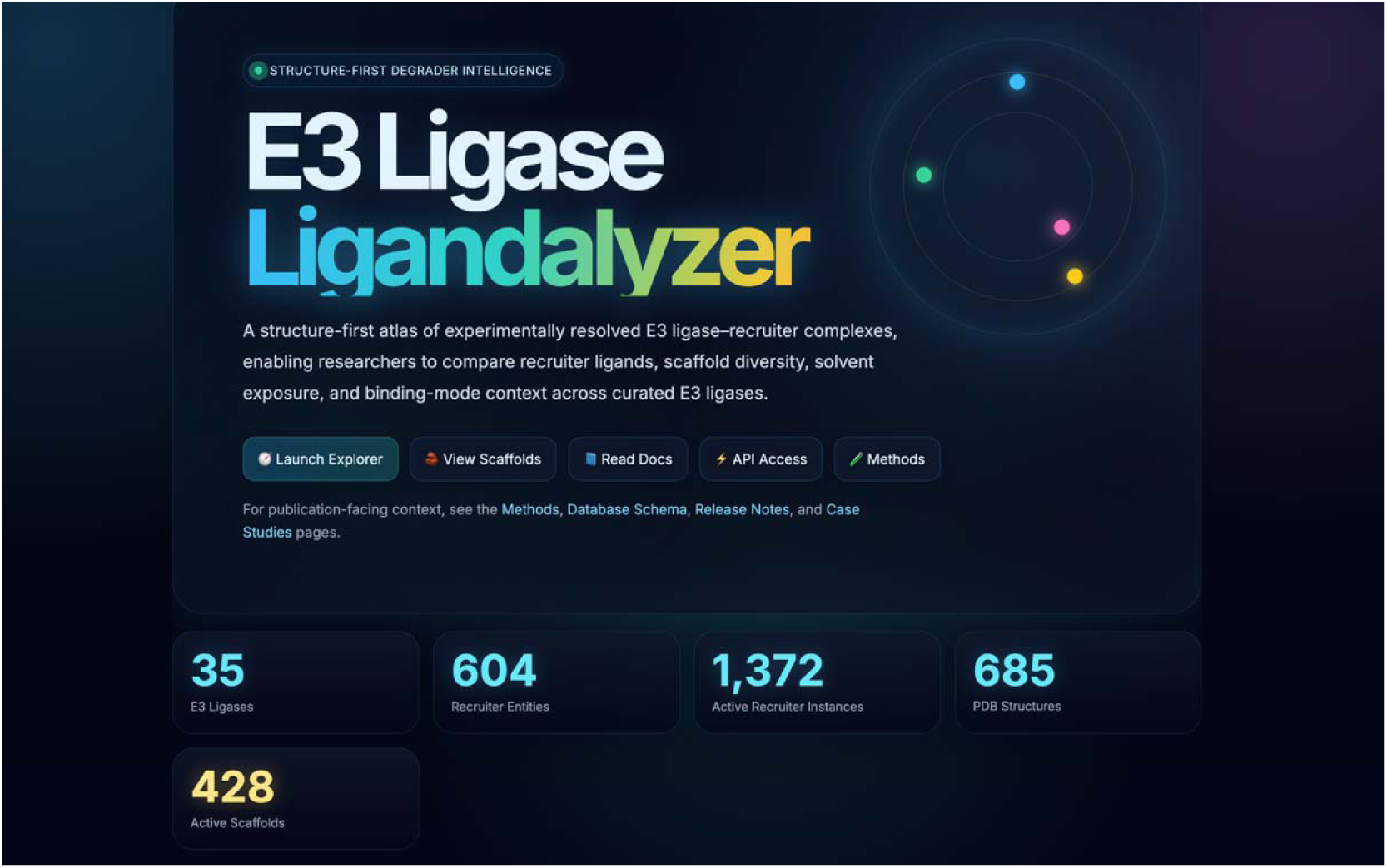
Production overview of E3 Ligase Ligandalyzer Version 1.0. The live interface summarizes the locked Version 1.0 dataset with 35 E3 ligases, 604 reconciled recruiter entities, 1,372 active structure-specific recruiter instances, 685 PDB structures, and 428 active scaffolds. Navigation links provide entry points to recruiter, ligase, scaffold, structural, and downstream-design modules. Alt text: Production E3 Ligase Ligandalyzer overview showing metric cards for 35 E3 ligases, 604 recruiter entities, 1,372 active recruiter instances, 685 PDB structures, and 428 active scaffolds.

**Figure 2.**
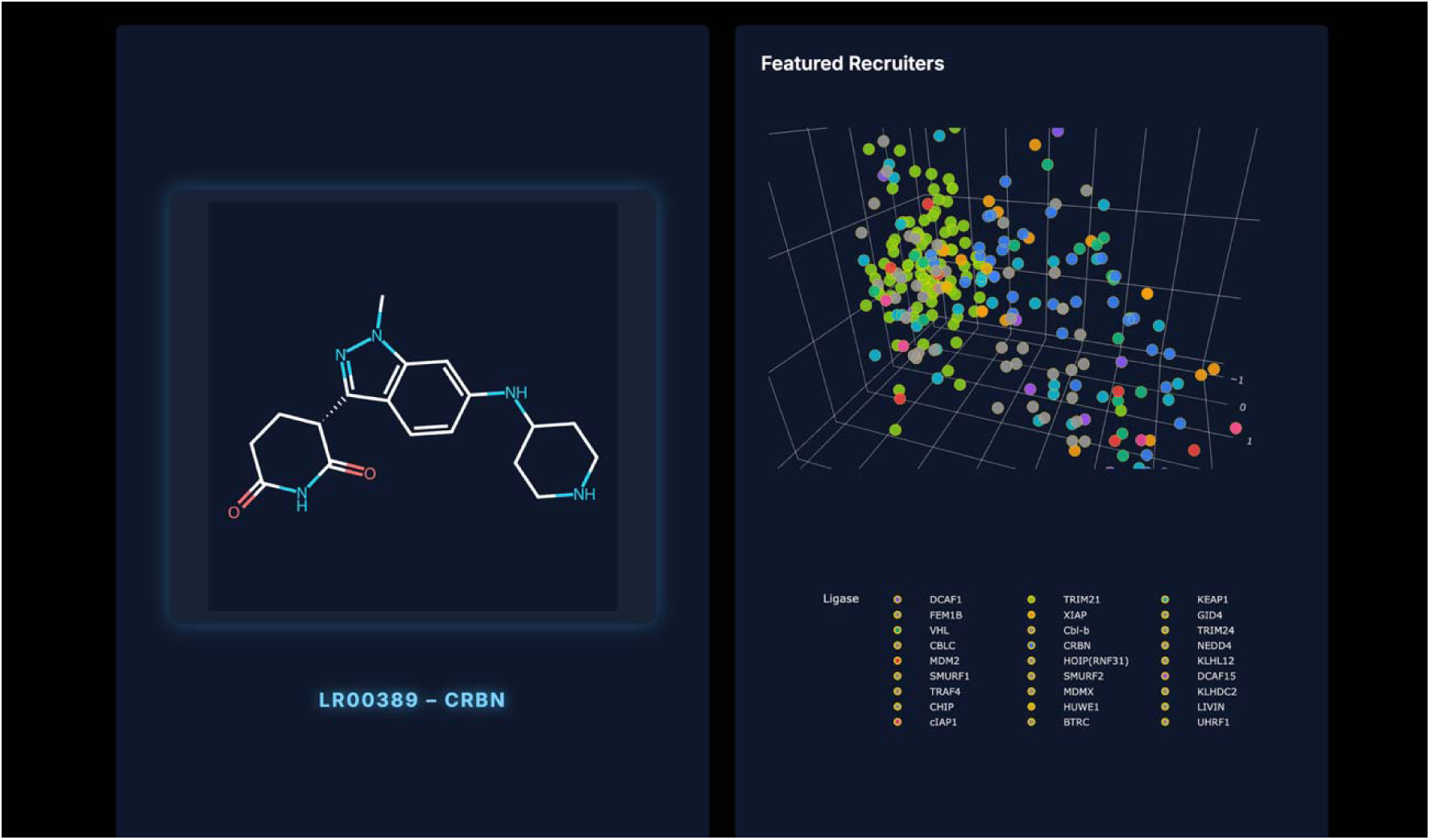
Interactive recruiter chemical-space view in the production E3 Ligase Ligandalyzer interface. The Featured Recruiters panel displays recruiter records in a three-dimensional view of calculated LogP, molecular weight, and QED; points are grouped by ligase association. Alt text: Production chemical-space interface showing a recruiter structure panel beside a three-dimensional Featured Recruiters plot of LogP, molecular weight, and QED grouped by ligase.

### Interactive overview of recruiter chemical space

The E3 Ligase Ligandalyzer homepage provides an interactive overview of recruiter chemical space, allowing users to explore entity-level physicochemical distributions across multiple dimensions. Reconciled recruiter entities are displayed in a three-dimensional scatter plot parameterized by quantitative estimate of drug-likeness (QED), calculated lipophilicity (LogP), and molecular weight (MW). Points are colored by ligase-recruiter association, enabling visual comparison of ligase-associated chemical regions and areas of overlap.

Hovering over a point reveals recruiter-entity metadata, including ligase association, scaffold class, physicochemical properties, and linked structural observations. Selecting a point opens a recruiter detail page with normalized two-dimensional chemistry, scaffold assignments, computed descriptors, and structure-specific solvent-exposure analyses. This interaction model supports movement from global entity-level chemical space to specific experimentally observed recruiter instances.

### Recruiter-level structural and solvent exposure analysis

Detailed recruiter pages present linked two-dimensional normalized chemistry and three-dimensional structure-specific representations (Figure 3). The two-dimensional depiction is annotated with atom-level SASA correspondence derived from Shrake-Rupley calculations, highlighting low, medium, and high solvent exposure in the observed instance. These annotations provide descriptive visual cues for positions that may merit follow-up evaluation.

**Figure 3.**
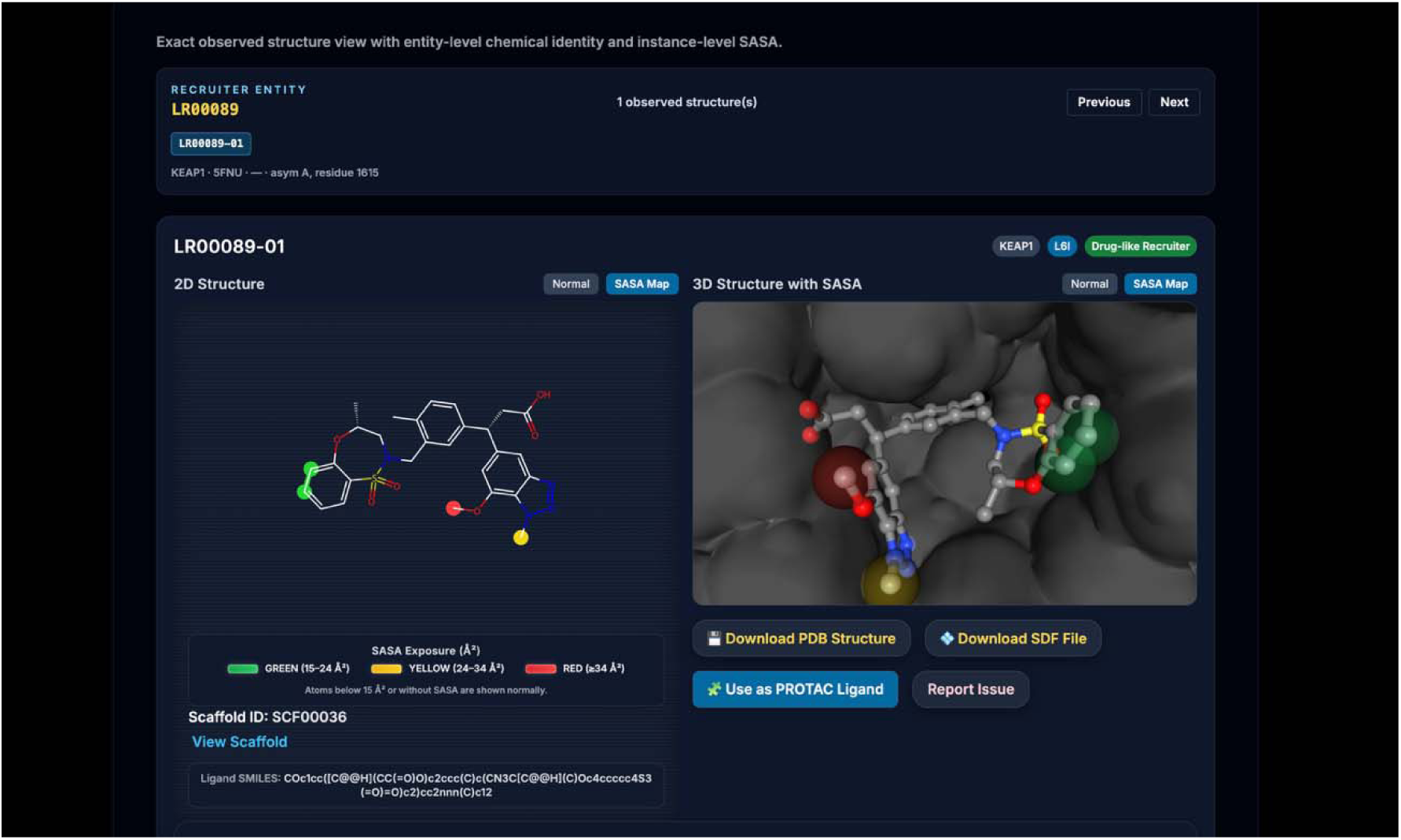
Recruiter-level 2D chemistry and observed 3D structural context for LR00089-01 (KEAP1, PDB 5FNU, L6I). The left panel shows normalized 2D recruiter chemistry with atom-level SASA correspondence. The right panel shows the experimentally observed PDB-coordinate recruiter pose within the KEAP1 pocket with SASA visualization. The observed 3D pose is not generated from the SDF. Alt text: Production recruiter detail view for LR00089-01 in KEAP1 PDB entry 5FNU, with normalized 2D chemistry and SASA correspondence on the left and the observed PDB-coordinate 3D ligand pose in its protein pocket on the right.

**Figure 4.**
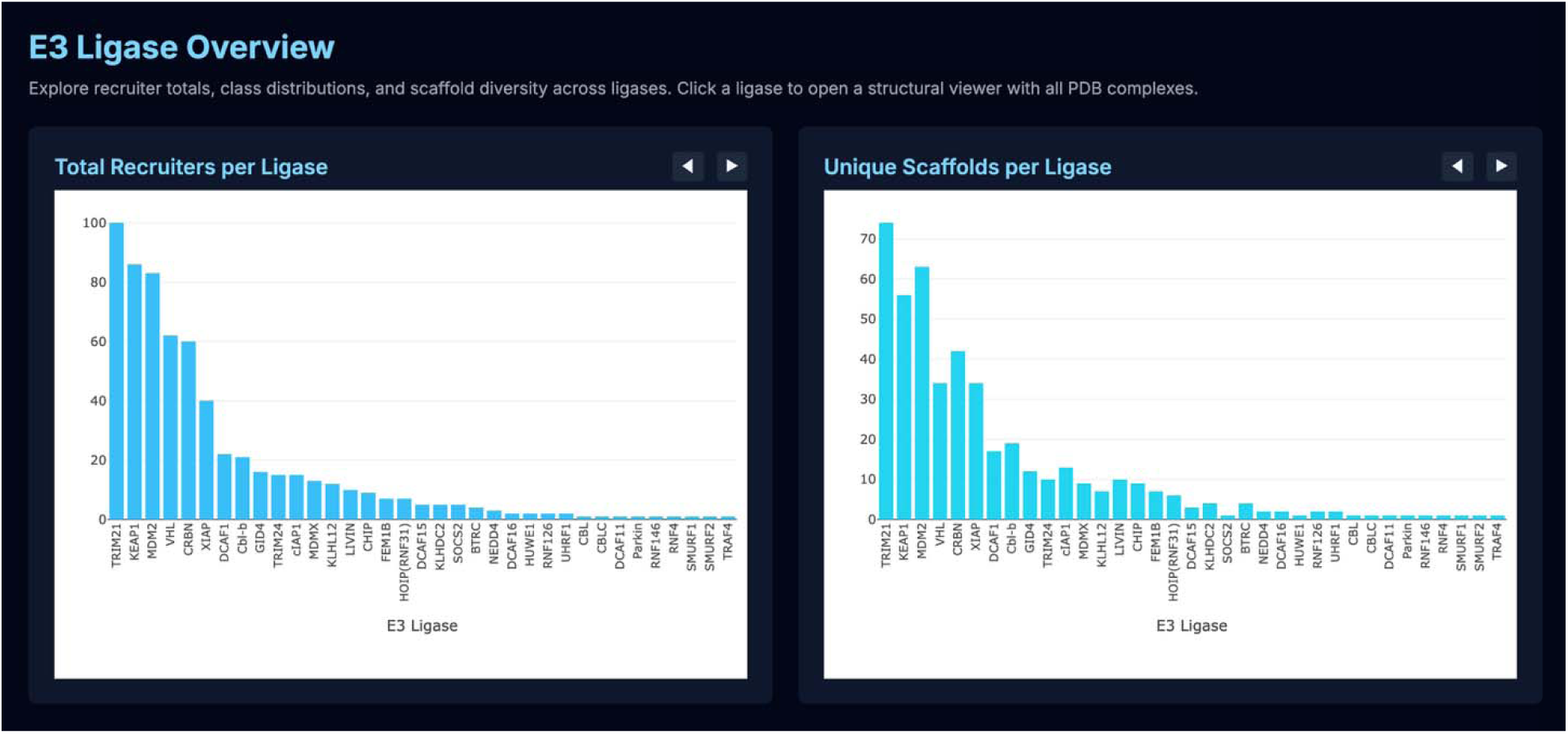
Ligase-centric exploration in E3 Ligase Ligandalyzer. The production Explorer displays recruiter totals and unique scaffold counts per ligase, enabling comparison of recruiter availability and scaffold coverage and direct access to ligase-specific structural exploration. Alt text: Production Ligase Explorer showing total recruiters per ligase and unique scaffolds per ligase.

The three-dimensional viewer displays the experimentally observed recruiter coordinates within the corresponding E3 ligase binding pocket, with ligand coloring reflecting SASA values. Users may toggle standard and SASA-mapped views, download structure files, or transfer normalized recruiter chemistry into PROTAC Builder. The two- and three-dimensional representations are linked through atom mapping, but the observed 3D pose is not generated from the SDF.

### Explorer module for recruiter selection by E3 ligase

E3 Ligase Ligandalyzer supports ligase-centric exploration through a module that summarizes recruiter-entity associations, active recruiter instances, scaffold diversity, and aggregated chemical properties for each E3 ligase represented in the database. In the locked Version 1.0 dataset, TRIM21, KEAP1, and MDM2 have the largest numbers of ligase-recruiter associations, while scaffold coverage remains heterogeneous across ligases. The explorer therefore distinguishes recruiter entities, active structural instances, and unique scaffolds rather than using the unqualified term recruiters for all three measures.

The ligase overview integrates association-level scaffold metrics, including scaffold diversity indices, normalized diversity scores, and recruiter-entity density. These measures provide a quantitative basis for comparing ligase-specific chemical space, while structure-specific instances retain the observed structural evidence underlying each association.

Selecting an individual ligase launches an aligned structure viewer that loads the available active recruiter instances associated with that ligase into a unified interactive scene (Figure 5).

**Figure 5.**
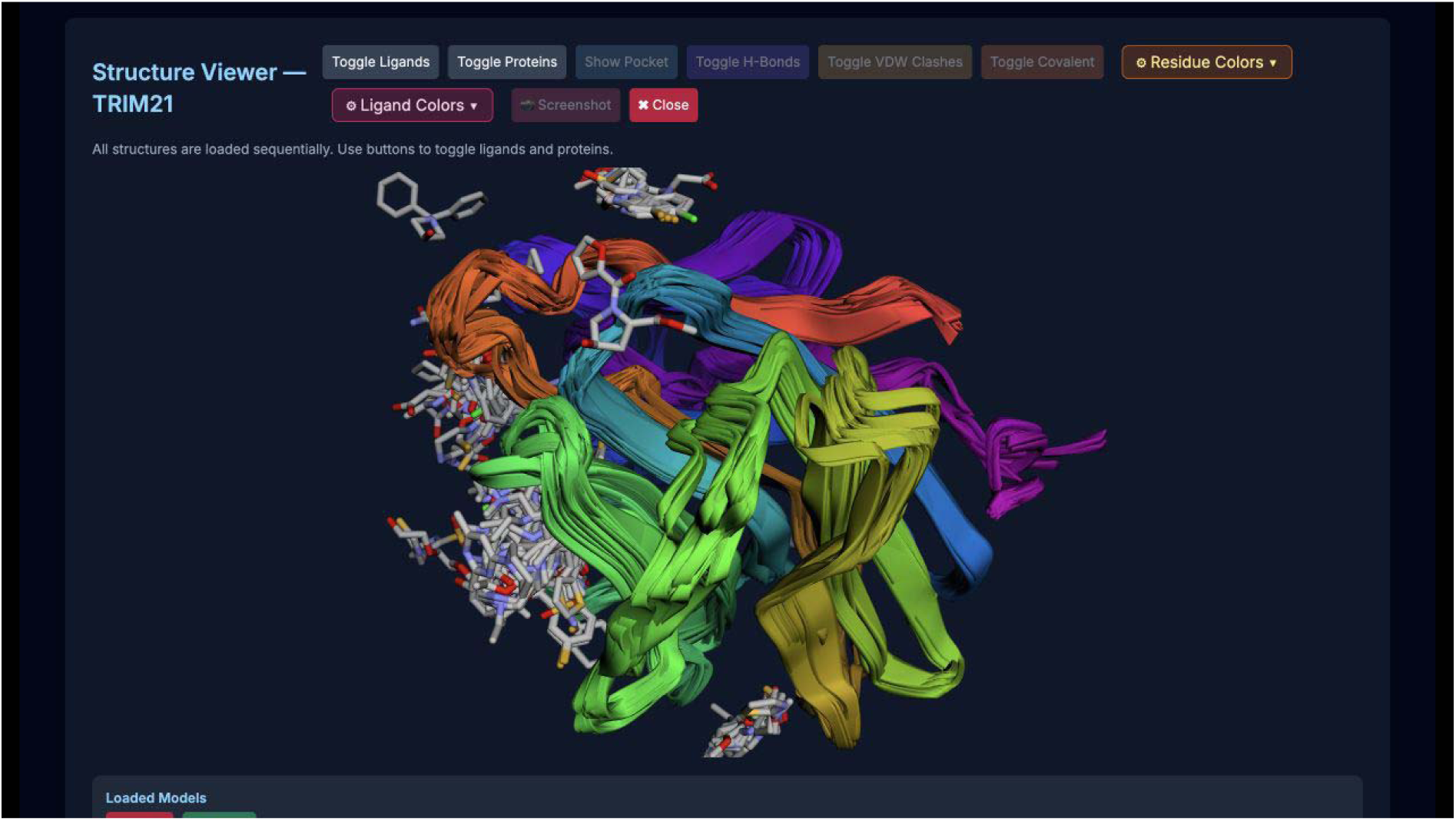
Aligned structure viewer for comparative analysis of active TRIM21 recruiter instances. The production viewer loads active recruiter instances sequentially and aligns their experimentally observed structures in a shared coordinate frame, enabling comparison of ligand orientations and binding-pocket context. Alt text: Production aligned structure viewer for TRIM21 showing active recruiter-bound structures in a shared coordinate frame with ligands and protein representations.

Structures are pre-aligned to a common reference frame, enabling visual comparison of observed binding modes, ligand orientations, and pocket geometries. Users may toggle individual recruiter instances or protein representations, selectively display binding pockets, and overlay interaction features. This view supports hypothesis generation around recruiter interchangeability, pocket adaptability, and ligase-specific design rules without treating the comparisons as predictions of productive degradation.

### Scaffold-centric analysis of recruiter chemical space

In addition to recruiter-entity and ligase-centered exploration, E3 Ligase Ligandalyzer provides a Scaffold Dashboard for scaffold-level analysis across profiled E3 ligases (Figure 6). The dashboard abstracts normalized recruiter-entity chemistry to Bemis-Murcko frameworks, allowing comparison of chemical architecture independent of substituent-level variation while retaining links to structural instances.

**Figure 6.**
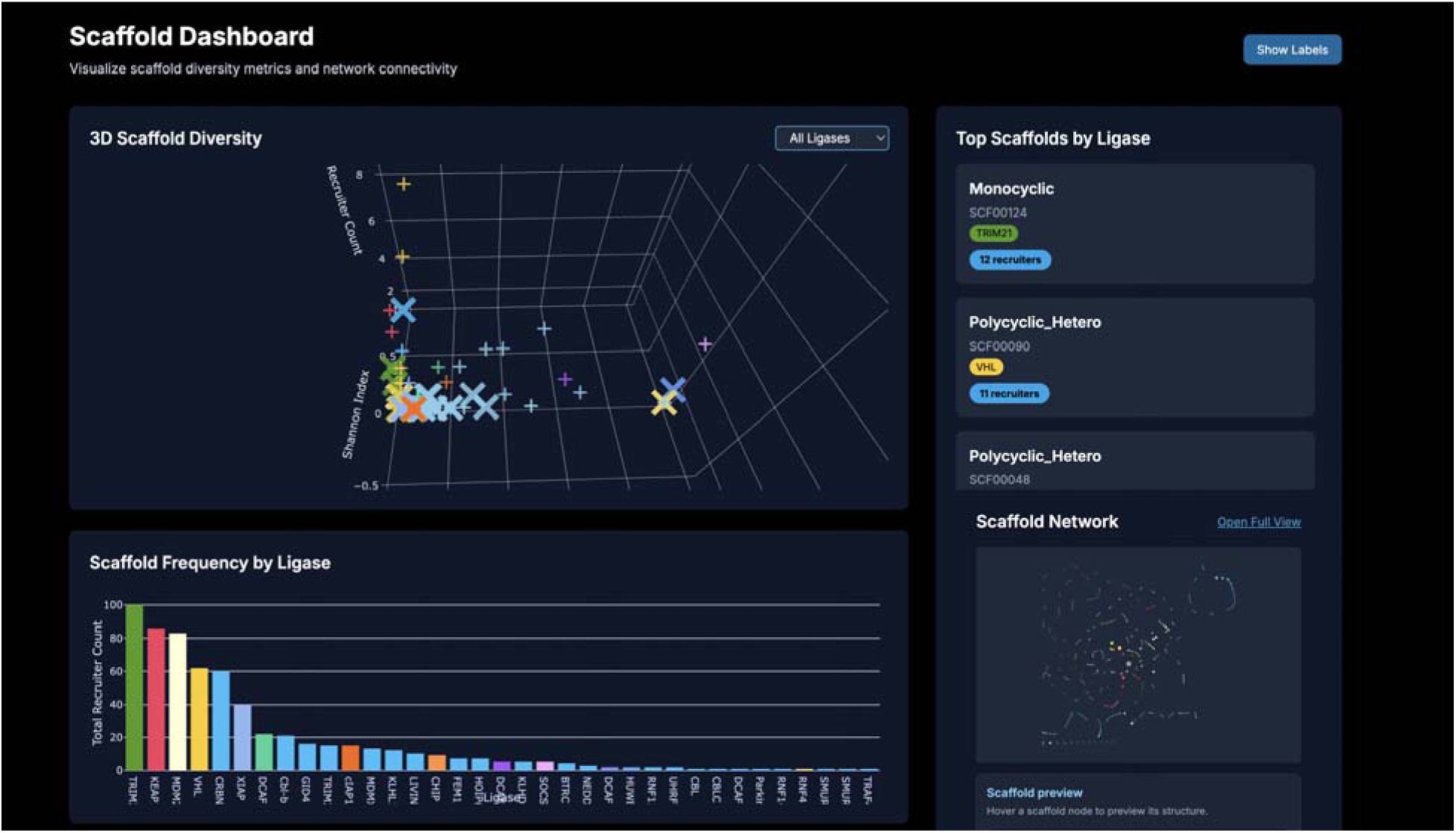
Scaffold-centric exploration in E3 Ligase Ligandalyzer. The production Scaffold Dashboard organizes active Version 1.0 recruiter chemistry by Bemis-Murcko framework and provides interactive three-dimensional diversity, scaffold-frequency, top-scaffold, and network views for examining scaffold usage across ligases. Alt text: Production Scaffold Dashboard showing an interactive three-dimensional scaffold-diversity plot, ligase selector, top scaffold cards, and scaffold network entry point.

As shown in Figure 6, scaffold diversity is visualized using a three-dimensional representation parameterized by recruiter density, Shannon diversity index, and total recruiter count. In this view, each point corresponds to a unique scaffold rather than an individual ligand, allowing users to distinguish broadly reused scaffolds from sparsely populated or ligase-specific scaffolds. This representation highlights patterns of scaffold reuse, diversification, and chemical convergence that are not apparent from ligand-level analysis alone. Scaffolds exhibiting high Shannon diversity and recruiter count indicate extensive chemical exploration within a shared core, whereas low-density scaffolds may represent emerging or underexplored chemotypes.

Complementary panels within the Scaffold Dashboard summarize scaffold frequency by ligase, providing a quantitative overview of how individual E3 ligases rely on distinct or overlapping scaffold classes (**Figure 6**). These distributions reveal marked heterogeneity in scaffold usage across ligases, reflecting differences in binding pocket architecture, historical medicinal chemistry efforts, and recruiter tractability. In parallel, a scaffold–ligase network visualization depicts connectivity between scaffolds and ligases, enabling identification of shared chemotypes spanning multiple E3 systems as well as ligase-restricted frameworks.

Together, the scaffold-centric analyses presented in Figure 6 support cross-ligase chemical reasoning by enabling users to evaluate recruiter space at the level of core molecular architecture. This abstraction facilitates identification of transferable scaffolds, prioritization of chemically versatile cores for degrader development, and recognition of ligase-specific design constraints that may limit scaffold reuse.

### Integration of expression-based filtering and tissue specificity

To provide biological context for recruiter and ligase selection, E3 Ligase Ligandalyzer integrates an expression-based filtering framework derived from the ELiAH E3 ligase expression atlas. The underlying dataset comprises expression profiles for 614 human E3 ligases across 2,292 tissue samples, spanning multiple physiologically relevant tissues, including skeletal muscle (366 samples), whole blood (291), thyroid (272), lung (225), heart atrial appendage (207), heart left ventricle (204), colon transverse (192), stomach (187), pancreas (183), and colon sigmoid (165). This broad coverage enables users to contextualize degrader design decisions across both broadly expressed ligases and those exhibiting tissue-enriched expression patterns.

Within E3 Ligase Ligandalyzer, ELiAH-derived expression data are exposed through a dedicated E3 Ligase Expression Atlas module that prioritizes interpretability and practical decision support. An overview dashboard summarizes dataset composition, tissue representation, and ligase coverage, while interactive gene and E3 ligase search functionality enables rapid querying of individual ligases across all available tissues (**Figure 7**). Rather than serving as a standalone transcriptomics resource, this module is designed to complement structure- and chemistry-driven analyses by providing immediate biological context during recruiter selection.

**Figure 7.**
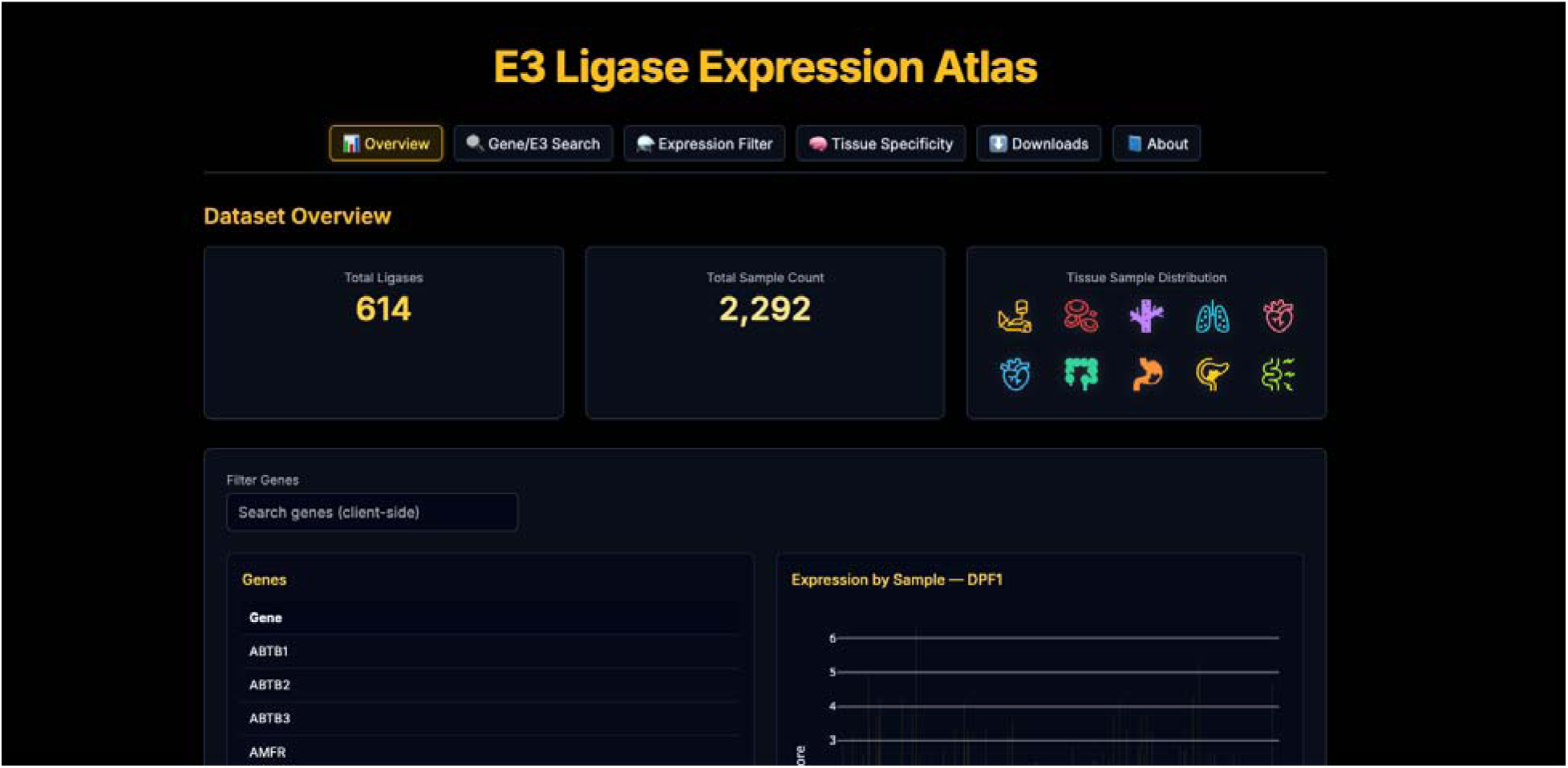
Overview of the E3 ligase expression dataset integrated into E3 Ligase Ligandalyzer. The E3 Ligase Expression Atlas module summarizes transcriptomic data for 614 human E3 ligases across 2,292 tissue samples derived from ELiAH. The overview dashboard reports total ligase coverage, sample counts, and tissue distribution, and provides an interactive gene and ligase search interface. Users may query individual E3 ligases to examine tissue-wise expression statistics, including mean Z-scores, enrichment values, and ligase specificity indices. This view provides global context for ligase expression patterns and supports rapid identification of ligases with broad or tissue-enriched expression profiles for downstream degrader design. *Alt text: E3 Ligase Expression Atlas overview summarizing expression data for 614 human E3 ligases across 2,292 tissue samples, with tissue coverage statistics and an interactive gene or ligase search interface*.

Expression data are presented as a contextual filter rather than a ranking or exclusion criterion. Users may examine ligase-level expression via a Gene/E3 Search interface that reports tissue-specific mean Z-scores, enrichment values, and a ligase specificity index, or via a Tissue Specificity view that visualizes expression distributions across selected tissues (Figure 8). These views allow users to quickly assess whether a ligase is ubiquitously expressed, moderately enriched, or relatively tissue-restricted, supporting informed prioritization decisions when multiple recruiter options are available.

**Figure 8.**
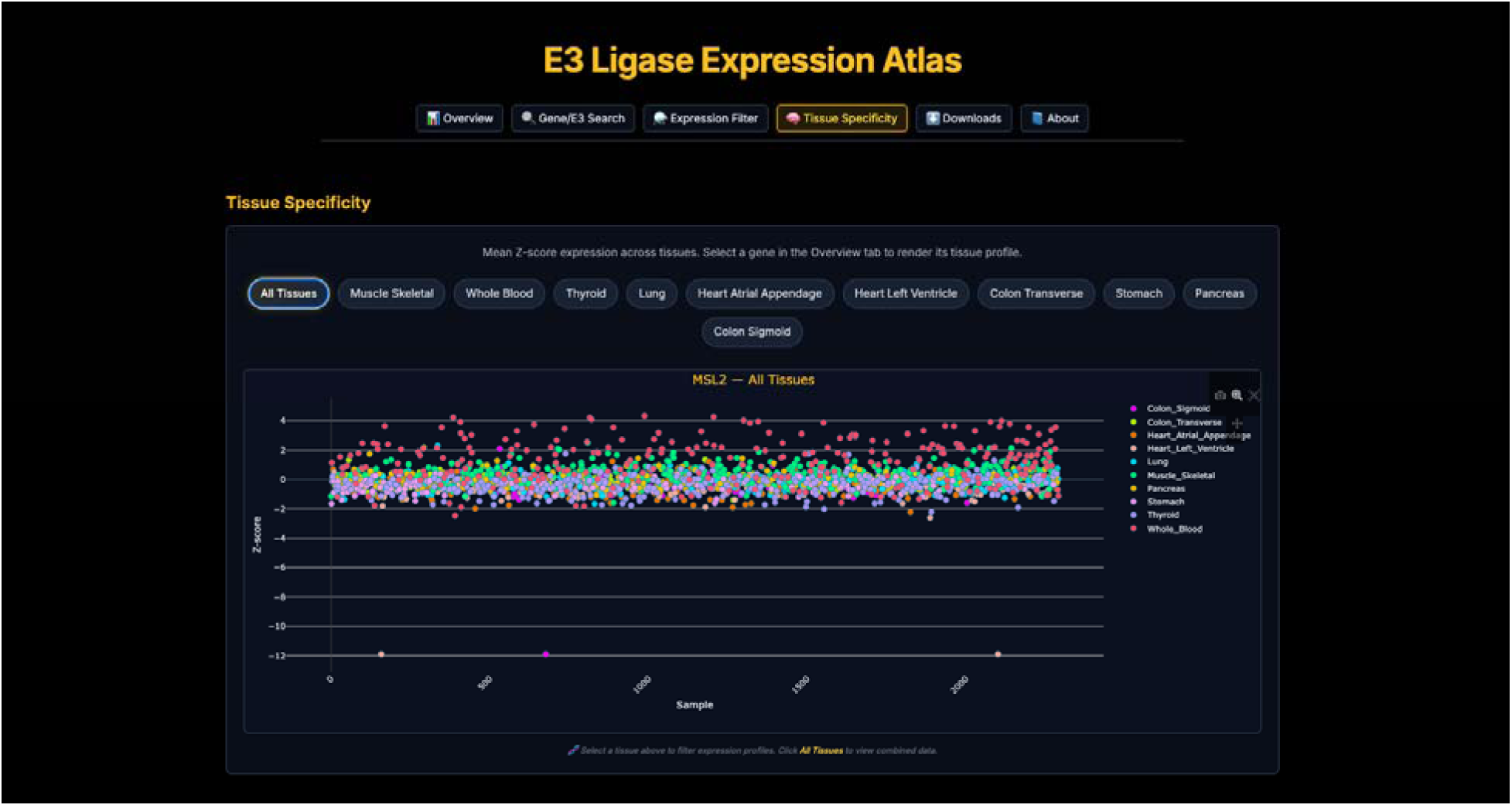
Tissue-specific expression visualization for individual E3 ligases within E3 Ligase Ligandalyzer. The Tissue Specificity view displays mean Z-score expression profiles of selected E3 ligases across individual tissues or aggregated datasets. Users can filter by tissue type to examine relative expression distributions and assess variability across samples. Expression profiles are visualized interactively, enabling comparison of tissue-enriched versus ubiquitously expressed ligases. This functionality is intended to provide biological context during recruiter and ligase selection, complementing structure- and chemistry-driven analyses without imposing expression-based ranking or exclusion criteria. *Alt text: Tissue-specific expression view displaying mean Z-score profiles for selected E3 ligases across individual tissues or aggregated datasets, with controls for comparing tissue-enriched and broadly expressed ligases*.

This functionality is particularly useful in early-stage degrader design, where expression considerations may influence ligase choice alongside structural compatibility, recruiter availability, and scaffold diversity. For example, users may deprioritize ligases with uniformly high expression across tissues when exploring strategies to improve selectivity, or, conversely, identify broadly expressed ligases when systemic degradation is desired. By enabling tissue filtering and side-by-side comparison of expression profiles, E3 Ligase Ligandalyzer lowers the barrier to incorporating biological relevance into structure-guided degrader workflows.

Importantly, E3 Ligase Ligandalyzer extends the usability of ELiAH expression data by integrating these views directly into the same environment used for recruiter exploration, scaffold analysis, and aligned structural inspection. Expression filters can be applied in parallel with ligase-centric recruiter statistics and scaffold diversity metrics, enabling a unified structure–chemistry–biology workflow without duplicating the functionality of expression-focused platforms. This integration ensures that biological context informs, rather than constrains, the selection of recruiters and ligases during targeted protein degradation design.

### User interaction and downstream workflow integration

E3 Ligase Ligandalyzer supports direct downstream integration with PROTAC Builder, enabling direct transition from recruiter evaluation to degrader construction. From recruiter-level views, users can select an individual recruiter ligand and initiate export to the PROTAC Builder environment through an attachment-site selection interface (Figure 9). This interface displays an atom-labeled molecular representation from which users select a specific attachment atom before export.

**Figure 9.** Interactive recruiter export from E3 Ligase Ligandalyzer to the PROTAC Builder environment. A recruiter ligand selected in E3 Ligase Ligandalyzer is prepared for downstream degrader design through an attachment-site selection interface. Users inspect an atom-labeled molecular representation, select a specific attachment atom, and export the recruiter to PROTAC Builder. Upon export, the recruiter is loaded as an editable two-dimensional chemical structure within an embedded sketch editor, enabling linker construction without manual file conversion. Alt text: Recruiter export workflow showing an atom-labeled molecular representation used to select an attachment atom, followed by transfer of the editable two-dimensional recruiter structure into PROTAC Builder for linker construction.

Upon transfer, recruiters are loaded into PROTAC Builder as fully editable two-dimensional chemical structures within an embedded sketch editor. This design enables immediate modification of recruiter chemistry, explicit definition of attachment points, and iterative linker construction without manual file conversion or intermediate data handling. By preserving recruiter identity and attachment-site metadata, this workflow facilitates continuity between recruiter selection and downstream constraint-driven degrader design.

This integration is designed to support geometry-aware, structure-first degrader development workflows, particularly those that rely on defined exit vectors and attachment constraints for subsequent modeling. Recruiters selected and prepared within E3 Ligase Ligandalyzer can therefore be directly incorporated into PROTAC Builder–based prototyping and, where applicable, propagated into constraint-driven ternary complex modeling pipelines.

Recent computational work has emphasized that nonstandard molecular representations and fragmented handoffs between modeling stages can limit reproducibility and pipeline-ready deployment in PROTAC design (35). The direct E3 Ligase Ligandalyzer–PROTAC Builder handoff provides a concrete implementation for addressing one upstream aspect of this problem: a recruiter can move from experimentally resolved structural inspection to editable degrader construction with an explicitly selected attachment site, without a separate manual file-conversion step.

Together, these features position E3 Ligase Ligandalyzer as an enabling upstream platform within a broader degrader design ecosystem, complementing expression-focused resources and degrader-centric databases while supporting rational, structure-informed decision-making at early stages of targeted protein degradation workflows.

## Discussion

Targeted protein degradation has matured into a structurally demanding modality in which recruiter selection, linker geometry, and ternary complex feasibility are tightly coupled (45). While recent advances in end-to-end structure prediction and generative modeling have accelerated exploratory design, reliable degrader development continues to depend on accurate representation of recruiter–ligase interactions and explicit consideration of geometric constraints. E3 Ligase Ligandalyzer was developed to address this need, enabling structure-first, constraint-aware decision-making at the earliest stages of degrader design.

A central premise underlying E3 Ligase Ligandalyzer is that experimentally determined recruiter–ligase complexes remain indispensable for geometry-driven workflows. Constraint-based docking frameworks, such as PRosettaC and related methods, rely on known anchor points, recruiter binding orientations, and linker exit vectors to generate physically plausible ternary complex ensembles. In contrast, although AlphaFold 3 has demonstrated impressive performance for many protein–protein and protein–ligand systems (46–48), it currently exhibits limitations in modeling degrader-specific geometries, particularly in cases involving flexible linkers, partial binding occupancy, or recruiter-induced conformational constraints (27). P4ward similarly illustrates how downstream automation still depends on well-specified upstream structural inputs: its ternary-complex workflow begins from binary protein-ligand complexes and PROTAC structures before docking, linker sampling, and scoring (37). As a result, accurate degrader modeling still benefits from curated structural inputs and explicit geometric constraints rather than fully unconstrained prediction.

These newer workflows clarify a practical role for a recruiter-centric structural resource. PCG explicitly preserves protein-bound recruiter and warhead geometries during linker sampling (38), DeepTernary adds an end-to-end SE(3)-equivariant prediction strategy (39), TERNIFY focuses the conformational search on the linker (40), and COMPASS evaluates linker candidates against ternary-complex formation and ubiquitination potential (41). Together with P4ward (37), these complementary approaches show how experimentally resolved recruiter-bound structures curated by E3 Ligase Ligandalyzer could serve as standardized E3-side starting inputs for multiple independent downstream modeling strategies rather than a single prescribed workflow.

E3 Ligase Ligandalyzer directly supports these approaches by aggregating and standardizing experimentally resolved E3 ligase-recruiter structures. The entity-instance architecture retains reconciled chemical identity alongside structure-specific observed coordinates and provenance. Solvent-exposure metrics, scaffold-level organization, and aligned pocket visualization enable users to examine structural context before engaging in linker design or ternary docking. The platform is an upstream comparative resource and does not rank ligases or predict productive degradation.

At the scale of the database, the API creates a testable route toward higher-throughput recruiter-side modeling. For a fixed POI–warhead input, a workflow could programmatically retrieve multiple experimentally resolved E3–recruiter complexes, apply consistent recruiter identifiers and structural coordinates, and repeat downstream ternary-complex modeling across alternative ligase–recruiter choices. We hypothesize that API-assisted setup will reduce manual structure preparation and improve input consistency relative to ad hoc PDB retrieval; this can be evaluated by comparing setup time, reproducibility, and model-generation success across the downstream workflows described above. Such scaling would increase the number of candidate PROTAC architectures that can be explored computationally while retaining experimentally grounded recruiter inputs, without treating structural feasibility as evidence of productive degradation.

Version 1.0 is a recruiter-focused upstream resource within a broader, interoperable degrader-design ecosystem. Its locked Version 1.0 dataset separates normalized recruiter chemistry from observed structural coordinates, allowing a recruiter entity and its documented structural instances to be retrieved reproducibly for downstream inspection. Direct integration with PROTAC Builder enables transfer of normalized chemistry for interactive degrader construction while retaining the distinction between chemistry representation and experimentally observed structural context.

E3 Ligase Ligandalyzer also complements warhead-focused tools within the same ecosystem. VLiSEMOD enables structure-guided analysis of viral protein targets for degrader development. Together, these tools reflect a modular design philosophy in which target selection, recruiter evaluation, and degrader geometry are treated as interconnected components of a unified, structure-first workflow.

The integration of expression-based filtering derived from ELiAH further enhances this framework by embedding biological context alongside structural and chemical considerations (21). Rather than imposing expression-based ranking or exclusion, E3 Ligase Ligandalyzer allows users to consider tissue expression profiles as an additional decision layer when evaluating recruiter–ligase combinations. This supports more informed prioritization without constraining exploration to a limited subset of well-characterized ligases.

In summary, E3 Ligase Ligandalyzer provides a focused, structure-centric platform for comparative analysis of E3 recruiter ligands and their associated ligases. By emphasizing experimentally grounded binding modes, scaffold relationships, and geometric constraints, and by integrating directly with downstream tools such as PROTAC Builder and VLiSEMOD, the platform supports rational, constraint-driven degrader design strategies that remain essential given current limitations in unconstrained modeling approaches. As targeted protein degradation continues to expand beyond canonical PROTAC architectures, tools that prioritize structural accuracy and workflow interoperability are likely to play an increasingly central role in the development of next-generation degradation therapeutics.

## Conclusion

Targeted protein degradation increasingly demands design strategies grounded in accurate molecular geometry, informed recruiter selection, and explicit consideration of linker constraints. E3 Ligase Ligandalyzer addresses a gap in recruiter-side comparison by organizing experimentally resolved E3 ligase-recruiter data around reconciled recruiter entities and their structure-specific instances. The locked Version 1.0 dataset provides 604 recruiter entities, 1,372 active recruiter instances, 685 PDB structures, and 35 E3 ligases for comparative chemical and structural exploration. Scaffold organization, physicochemical characterization, solvent exposure metrics, and aligned structural visualization support hypothesis generation without claiming predictive ranking or validated linker tolerance.

Rather than functioning as an isolated database, E3 Ligase Ligandalyzer is designed to operate as an upstream decision-support layer within a broader ecosystem of structure-informed tools. Direct integration with PROTAC Builder enables an immediate transition from recruiter selection to degrader construction, while interoperability with complementary platforms, such as VLiSEMOD, supports coherent target–recruiter–linker design workflows. This modular, structure-first philosophy reflects the growing recognition that accurate degrader modeling depends on the quality and interpretability of its foundational inputs.

As the landscape of targeted protein degradation continues to expand beyond a small subset of canonical E3 ligases into increasingly diverse therapeutic contexts, resources that enable rational, recruiter-centric comparisons across ligases will become increasingly important. By lowering the barrier to structure-informed recruiter evaluation and embedding this capability within an extensible computational ecosystem, E3 Ligase Ligandalyzer provides a foundation for more systematic, geometry-aware design of next-generation degradation therapeutics.

## Data Availability

E3 Ligase Ligandalyzer Version 1.0 is freely accessible without registration at https://e3ligandalyzer.com. The curated recruiter-entity, recruiter-instance, ligase, scaffold, structural, and calculated descriptor data can be queried and downloaded through the public API documentation at https://e3ligandalyzer.com/api-reference. Version 1.0 table exports, including recruiter, recruiter-instance, scaffold, descriptor, and SASA data, are available through the public download index at https://e3ligandalyzer.com/api/download/tables. Dataset-generation scripts and supporting computational workflows are available at https://github.com/schurerlab/E3-Ligandalyzer-Scripts. Source structural data were obtained from the RCSB Protein Data Bank, and PDB accession identifiers and structure-specific provenance are retained in the database records.

All quantitative results reported in this article refer to the locked E3 Ligase Ligandalyzer Version 1.0 dataset with a database cutoff of 8 September 2026. The associated SQLite artifact has SHA-256 a7897716e7db29adb8569a58de0b5b3819b89f5c9cc0a509437b33e64852d7a3. The live platform may evolve in later releases, but manuscript counts remain reproducible against the locked Version 1.0 dataset.

## Funding

This work was supported by the Florida Department of Health [grant number 23B16]; the Congressionally Directed Medical Research Programs [award number 1420842]; and the U.S. National Institutes of Health, including the National Cancer Institute [grant number P30 CA240139] and the National Library of Medicine [grant numbers U01 LM012630 and R01 LM013391]. The funding agencies had no role in the design of the study; the collection, analysis, or interpretation of the data; the preparation of the manuscript; or the decision to submit the article for publication.

## Acknowledgements

We thank the Sylvester Comprehensive Cancer Center and the Department of Molecular and Cellular Pharmacology at the University of Miami for sustained institutional support. We are grateful to the Frost Institute for Data Science & Computing for access to high-performance computing infrastructure that supported the large-scale structural analyses and data-processing workflows underlying this resource. We also acknowledge the RCSB Protein Data Bank and the developers and maintainers of the open-source computational chemistry, cheminformatics, and structural biology software used in the development of E3 Ligase Ligandalyzer.

## Conflict of Interest

The authors declare no competing interests.

## Notes

https://e3ligandalyzer.com

https://e3ligandalyzer.com/api-reference

https://e3ligandalyzer.com/api/download/tables

https://github.com/schurerlab/E3-Ligandalyzer-Scripts

### Competing Interest Statement

The authors have declared no competing interest.

